# CD8 T lymphocytes infiltrate the kidneys and correlate with disease progression in B6.*NZMSle1/Sle2/Sle3* lupus mice

**DOI:** 10.1101/2025.11.03.685164

**Authors:** Pauline Montigny, Selda Aydin, Anne-Lise Maudoux, Nicolas Van Baren, Davide Brusa, Nicolas Dauguet, Frédéric Houssiau, Bernard Lauwerys, Pierre G. Coulie

## Abstract

**Background:** Lupus nephritis (LN) is a severe manifestation of systemic lupus erythematosus (SLE) characterized by immune-mediated renal damage. While intrarenal CD8^+^ T cell infiltration has been linked to disease activity in patients, their pathogenic contribution remains unclear, partly due to the lack of mechanistic insight from murine models.

**Methods:** We investigated renal CD8^+^ T cell infiltration in female B6.NZMSle1/Sle2/Sle3 lupus-prone mice across different ages, comparing them to C57BL/6 controls. Histopathology was assessed using human-derived NIH Activity and Chronicity Indices, complemented by fibrosis quantification, immunohistochemistry, and digital image analysis. Kidney T cell subsets were evaluated by flow cytometry, and transcriptomic profiling was performed using microarrays.

**Results:** Lupus-prone mice developed progressive kidney injury resembling human LN, with increasing NIH activity scores, collagen deposition, however with interindividual heterogeneity. CD8^+^ T cell infiltrates were significantly elevated in lupus kidneys as early as 3 months, rising with age and correlating with histological activity. CD8^+^ T cells localized to periglomerular and peritubular regions but did not predominate over CD4^+^ T cells, as confirmed by flow cytometry. Transcriptomic analyses revealed age-dependent upregulation of interferon (IFN)-stimulated genes, B cell–associated transcripts, and extracellular matrix remodeling pathways, while T cell– related signatures were more variable.

**Conclusions:** B6.NZMSle1/Sle2/Sle3 mice recapitulate several histopathological and molecular features of human LN, including progressive fibrosis and intrarenal CD8^+^ T cell infiltration that correlate with disease severity. However, the absence of CD8^+^ predominance suggests limitations of this model for dissecting CD8^+^ T cell-specific contributions to LN pathogenesis. Yet, these findings underscore the need to identify renal antigens driving CD8^+^ T cell responses in human LN.

## Introduction

Systemic lupus erythematosus (SLE) is a chronic systemic autoimmune disease that predominantly affects women of childbearing age and presents with a broad spectrum of clinical manifestations. Lupus nephritis (LN) is the most common severe manifestations of SLE: despite multimodal immunosuppressive therapy, 10–30% of patients progress to end-stage renal disease [1]. Current guidelines recommend initial therapy with low-dose cyclophosphamide or mycophenolate mofetil (MMF) in combination with glucocorticoids (GC), followed by maintenance therapy with MMF or azathioprine and progressive tapering of GC [2]. More recently, anti-BLyS belimumab and voclosporin (a new calcineurin inhibitor) have been approved for the treatment of LN, and anti-CD20 obinutuzumab might follow soon. Whether these targeted therapies should be given to all patients is debated but is advised by both the ACR guidelines [3] and the EULAR recommendations [4].

The pathophysiology of LN is not fully understood. Disease onset appears to involve glomerular deposition of immune complexes containing nuclear material, leading to glomerulonephritis. B lymphocytes activated by nucleic acids, with CD4^+^ T cell help, produce high-affinity autoantibodies against DNA and other nuclear antigens. The breakdown of tolerance to nuclear components is most likely linked to defective clearance mechanisms, resulting in the accumulation of immunogenic nuclear debris.

In addition to glomerular injury, kidneys from LN patients show tubulointerstitial damage with immune cell infiltrates that include T lymphocytes, among them CD8^+^ T cells. Only a few studies have systematically compared CD4^+^ and CD8^+^ T cell subsets in LN kidney tissue using immunohistochemistry. In one study analyzing 26 kidneys, CD8^+^ T cells predominated [5], while another study of 18 kidneys reported comparable proportions of CD4^+^ and CD8^+^ cells [6]. A study from 1990 quantified CD4^+^ and CD8^+^ T cells in the renal interstitium of 35 LN patients receiving immunosuppressive therapy, correlating these counts with estimated glomerular filtration rate (eGFR) at biopsy. CD4^+^ T cells predominated in 19 cases, whereas CD8^+^ cells predominated in 16 [7]. More recently, in 25 LN kidneys, CD8^+^ T cells were more abundant than CD4^+^ T cells, CD20+ B cells, or CD68^+^ macrophages, and their infiltration correlated with the NIH Activity Index [8]. In 2012, semiquantitative analyses of 25 LN biopsies (including repeated samples) found CD4^+^ T cells to predominate in two-thirds of cases, although small periglomerular infiltrates were enriched in CD8^+^ T cells. TCR sequencing of these microdissected clusters revealed oligoclonality, with some clones persisting in later biopsies and detectable in autologous peripheral blood CD8^+^ T cells [9]. Our group applied digital immunohistochemistry to 37 proliferative LN biopsies and found that CD3^+^, CD8^+^, and CD20^+^ cell infiltrates correlated with reduced eGFR at biopsy, with the strongest statistical association observed for CD8^+^ cells [10]. Furthermore, in 40 LN biopsies, we identified senescent p16-positive cells significantly associated with reduced eGFR both at biopsy and at 5-year follow-up. Notably, glomerular p16-positive cells showed marked spatial co-distribution with periglomerular CD8^+^ T cells [11].

Taken together, these findings support a role for CD8^+^ T lymphocytes in LN pathogenesis. However, in the absence of identified kidney-specific antigens recognized by these CD8^+^ T cells, bystander recruitment cannot be excluded—that is, CD8^+^ T cells may recognize antigens unrelated to the kidney and be attracted secondarily by chemokines secreted locally by inflammatory cells.

Mouse models of LN provide further insights into CD8^+^ T cell involvement. In these models, lupus either arises spontaneously, as in MRL/lpr, (NZW/BXSB)F1, (NZB/NZW)F1, or NZM2410 mice, or is experimentally induced. In the NZM2410 strain, Wakeland and colleagues identified three recessive loci (Sle1, Sle2, Sle3) associated with lupus susceptibility and individually transferred onto a C57BL/6 background [12]. Later, the same group generated the triple congenic B6.NZMSle1/Sle2/Sle3 strain, which develops severe systemic autoimmunity and uniformly fatal glomerulonephritis [13]. Interestingly, we also observed glomerular p16^+^ cells in B6.NZMSle1/Sle2/Sle3 kidneys, similar to those detected in human LN [14].

Here, we quantitated and characterized CD8^+^ T cell infiltrates in the kidneys of B6.NZMSle1/Sle2/Sle3 mice across different ages.

## Materials and Methods

### Mice

B6.NZMSle1/Sle2/Sle3 mice were kindly provided by Prof. Edward K. Wakeland (Center for Immunology, University of Texas Southwestern Medical Center, Dallas) and maintained alongside healthy C57BL/6 mice in a Specific Pathogen-Free (SPF) facility, in compliance with national ethical regulations. Mice were euthanized at different ages for kidney and spleen collection.

### Histological scoring, immunohistochemistry, and quantitative image analysis

Eighteen healthy female C57BL/6 mice and thirty-two lupus-prone female B6.NZMSle1/Sle2/Sle3 mice were included in the histopathological analyses. Kidneys were dissected longitudinally, formalin-fixed, and paraffin-embedded. Sections were stained with Hematoxylin and Eosin (H&E) and Periodic Acid–Schiff (PAS) following manufacturer’s protocols. CD8^+^ T cells were detected using a monoclonal anti-CD8α rabbit antibody (Cell Signaling Technology, #98941, 1:200) and an HRP-based secondary detection system. Collagen types I and III were stained with Sirius Red (ThermoFisher, #B21693). Quantification of CD8^+^ cells and collagen fibers was performed using the HALO image analysis software. The National Institutes of Health Activity Index (NIH AI) and Chronicity Index (NIH CI) were scored by a single blinded pathologist [15].

### Flow cytometry

Kidneys from 32 lupus-prone B6.NZMSle1/Sle2/Sle3 mice and 5 healthy C57BL/6 controls were processed for flow cytometry. A single kidney per mouse was minced into small fragments and mechanically dissociated in RPMI 1640 medium. Cell suspensions were filtered through a 70 µm mesh, centrifuged, and resuspended in PBS-EDTA. To block nonspecific antibody binding, cells were pre-incubated with Normal Goat Serum (NGS) before staining with anti-CD8 (BD Biosciences, #552877), anti-CD3 (BD Biosciences, #562286), and 7-AAD (BD Biosciences, #559925). Spleens were mechanically dissociated in RPMI 1640 medium and filtered through a 70 µm mesh. Mononuclear cells were isolated by density-gradient centrifugation. After blocking with NGS, cells were stained with the same antibody panel. Data acquisition was performed on a BD FACSAria flow cytometer.

### Microarrays

RNA was extracted from mouse kidneys using TriPure Isolation Reagent (Roche Life Science) according to the manufacturer’s instructions and hybridized to Affymetrix GeneChip Mouse Gene 2.0 ST arrays. Raw CEL files were processed in Qlucore Omics Explorer v3.10 using RMA sketch normalization. Gene expression levels were compared between kidneys from 3-month-old mice (controls) and older mice using a two-tailed unpaired t-test with batch correction. A q-value (false discovery rate) threshold of 0.01 identified 72 differentially expressed genes. Gene Set Enrichment Analysis (GSEA) was performed using default parameters, with t-test statistics as the ranking metric. The *Ifna*-induced, *Ifng*-induced, and *Spermatogenesis* gene sets were retrieved from the murine MSigDB Hallmark collection [16]. T-cell– and B-cell–associated gene signatures were manually curated from various sources and are listed hereunder: T-cell–associated genes: *Cd2, Cd3d, Cd3g, Cd4, Cd6, Cd8a, Cd8b1, Cd28, Cd40lg, Cd247, Csf2, Cst7, Ctsw, Cxcr3, Eomes, Foxp3, Fyb, Gata3, Gimap5, Gzmb, Gzmc, Icos, Ifng, Il2, Il2ra, Il2rb, Il2rg, Il4, Il5, Il9, Il13, Il17a, Il17f, Il7r, Il21, Il22, Lag3, Lat, Lck, Lef1, Lta, Ltb, Mal, Pdcd1, Prf1, Runx2, Sh2d1a, Sema7a, Sh2d1a, Stat4, Tbx21, Tcf7, Tnfsf11, Trac, Trbc2, Zap70;* B-cell–associated genes: *Blk, Blnk, Btk, Cd19, Cd22, Cd38, Cd72, Cd79a, Cd79b, Cd200, Cebpb, Gcsam, H2DMa, Ighg3, Igll1, Ighm, Iglc2, Iglc3, Irf8, Jchain, Lyn, Ms4a1, Pax5, Pou2af1, Prdm1, Spib, Stat6, Syk, Tnfsf13b, Tnfrsf13b, Tnfrsf13c, Tnfrsf1*.

## Results

### Progression of lupus nephritis with age in B6/Sle1.Sle2.Sle3 mice

Kidneys and spleens were collected from female B6.Sle1.Sle2.Sle3 mice aged 3, 6, and 9–14 months, as well as from age-matched healthy C57BL/6 controls. Histological progression of lupus nephritis was assessed using the NIH Activity Index (NIH AI, range 0-24) and NIH Chronicity Index (NIH CI, range 0-12). The NIH AI reflects active lesions of nephritis, whereas the NIH CI captures irreversible damage, mainly fibrotic and atrophic changes. Each index is based on a set of parameters scored from 0 (absent) to 3 (severe)[17]. NIH AI parameters include glomerular hypercellularity, leukocyte exudation, karyorrhexis/fibrinoid necrosis, cellular crescents, hyaline deposits, and interstitial inflammation. NIH CI parameters include glomerulosclerosis, fibrous crescents, tubular atrophy, and interstitial fibrosis. Although these indices were originally developed for human lupus nephritis, they had not yet been applied to the B6.Sle1.Sle2.Sle3 model.

As shown in Fig. 1A (left panel), NIH AI scores in lupus-prone mice were already elevated at 3 months compared with C57BL/6 controls. Scores increased with age, reaching a median of 6.0 in 9–14-month-old mice. This range is comparable to values reported in patients with newly diagnosed lupus nephritis, where a median NIH AI score of 8.5 has been described [18]. From 6 to 14 months of age, lupus mice displayed considerable interindividual variability in NIH AI scores, mirroring the heterogeneity observed in patients. In contrast, NIH CI scores provided limited discrimination between healthy and lupus mice, or between younger and older lupus mice (Fig. 1A, right panel).

**Fig. 1.**
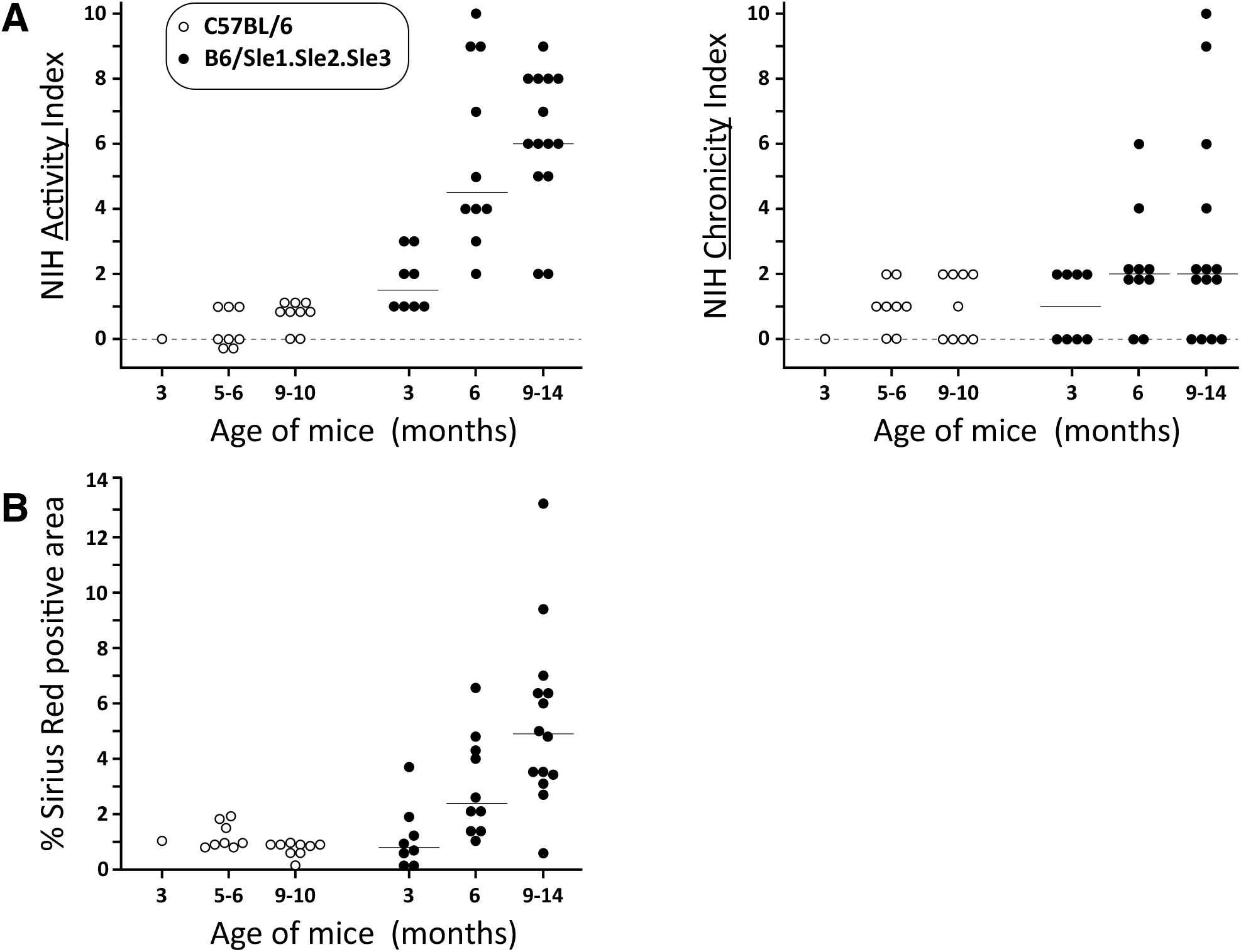
Histological evaluation of kidney disease severity in B6/Sle1.Sle2.Sle3 mice. **(A)** NIH Activity and Chronicity Indices in lupus-prone and healthy mice according to age. Kidneys from mice of different ages were processed using the same protocols as for human renal biopsies. Consecutive formalin-fixed, paraffin-embedded (FFPE) sections were stained with H&E or PAS and scored by an expert nephropathologist. Results are grouped by age category; horizontal lines indicate median values. **(B)** Quantification of kidney fibrosis by Sirius Red staining. FFPE kidney sections were stained with Sirius Red and digitalized. Collagen deposition was quantified within the renal parenchyma, excluding capsular and hilar regions, using the HALO image analysis platform. Results are expressed as the percentage of stained area relative to the analyzed parenchymal surface; horizontal lines indicate median values.

To assess kidney fibrosis—a hallmark of chronic damage in human lupus nephritis—we stained collagen types I and III with Sirius Red, followed by digital quantification of stained areas. As shown in Fig. 1B, fibrosis was already detectable in lupus mice older than 3 months and increased with age, reaching a mean of 6.2% of kidney parenchyma stained. Comparable values have been reported in patient biopsies [19].

In summary, B6.Sle1.Sle2.Sle3 mice develop progressive kidney disease characterized by histological features and grading patterns that closely resemble human LN. These findings extend the histopathological characterization of this model beyond the original description by Wakeland’s group.

### Progression of CD8^+^ T cell infiltration in the kidneys of B6/Sle1.Sle2.Sle3 mice

We quantified renal CD8^+^ T cell infiltration by digital immunohistochemistry on longitudinal whole-kidney sections from B6.Sle1.Sle2.Sle3 mice and age-matched healthy controls. In healthy mice, CD8^+^ cells were rare, representing <0.3% of all nucleated cells, with a median of 0.08%, and showed no increase with age (Fig. 2A). In contrast, lupus-prone mice exhibited significantly higher proportions of CD8^+^ cells as early as 3 months of age (median 0.57%, *p* < 0.0001), with all values exceeding those of healthy controls. CD8^+^ cell infiltration increased progressively with age, reaching median values of 1.56% at 6 months and 2.81% at 9–14 months (Fig. 2A).

**Fig. 2.**
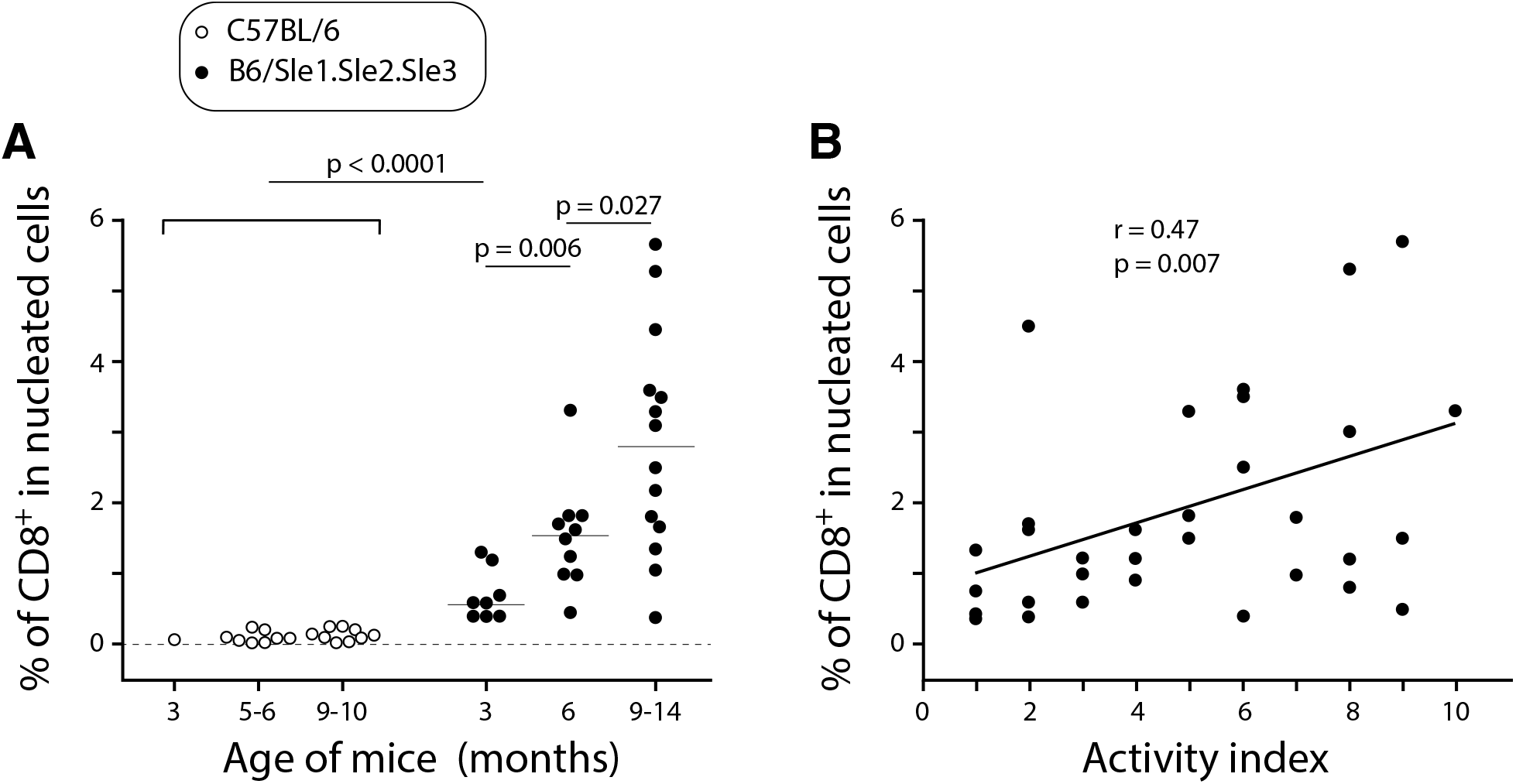
Digital quantification of renal CD8^+^ T cell infiltration in healthy and B6.Sle1.Sle2.Sle3 mice. **(A)** Proportions of CD8^+^ cells among all nucleated cells, quantified on FFPE kidney sections stained with an anti-CD8α monoclonal antibody and counterstained with hematoxylin. **(B)** Pearson correlation between CD8^+^ cell infiltration and NIH Activity Index scores in kidneys from lupus-prone mice.

In lupus mice, the proportions of CD8^+^ infiltrating cells correlated positively with histological disease activity scores (Fig. 2B), in line with findings previously reported in patients with lupus nephritis by Couzi and colleagues [8].

CD8^+^ T cells were distributed throughout the renal parenchyma without a uniform localization pattern. They were frequently detected in peritubular (Fig. 3A, C) and periglomerular areas (Fig. 3A, B). Only some glomeruli were surrounded by multiple CD8^+^ T cells (Fig. 3B). These periglomerular infiltrates appeared more common in proximity to perivascular mononuclear cell aggregates (Fig. 3D-E). CD8^+^ T cells were also detected diffusely within these inflammatory infiltrates (Fig. 3D, E).

**Fig. 3.**
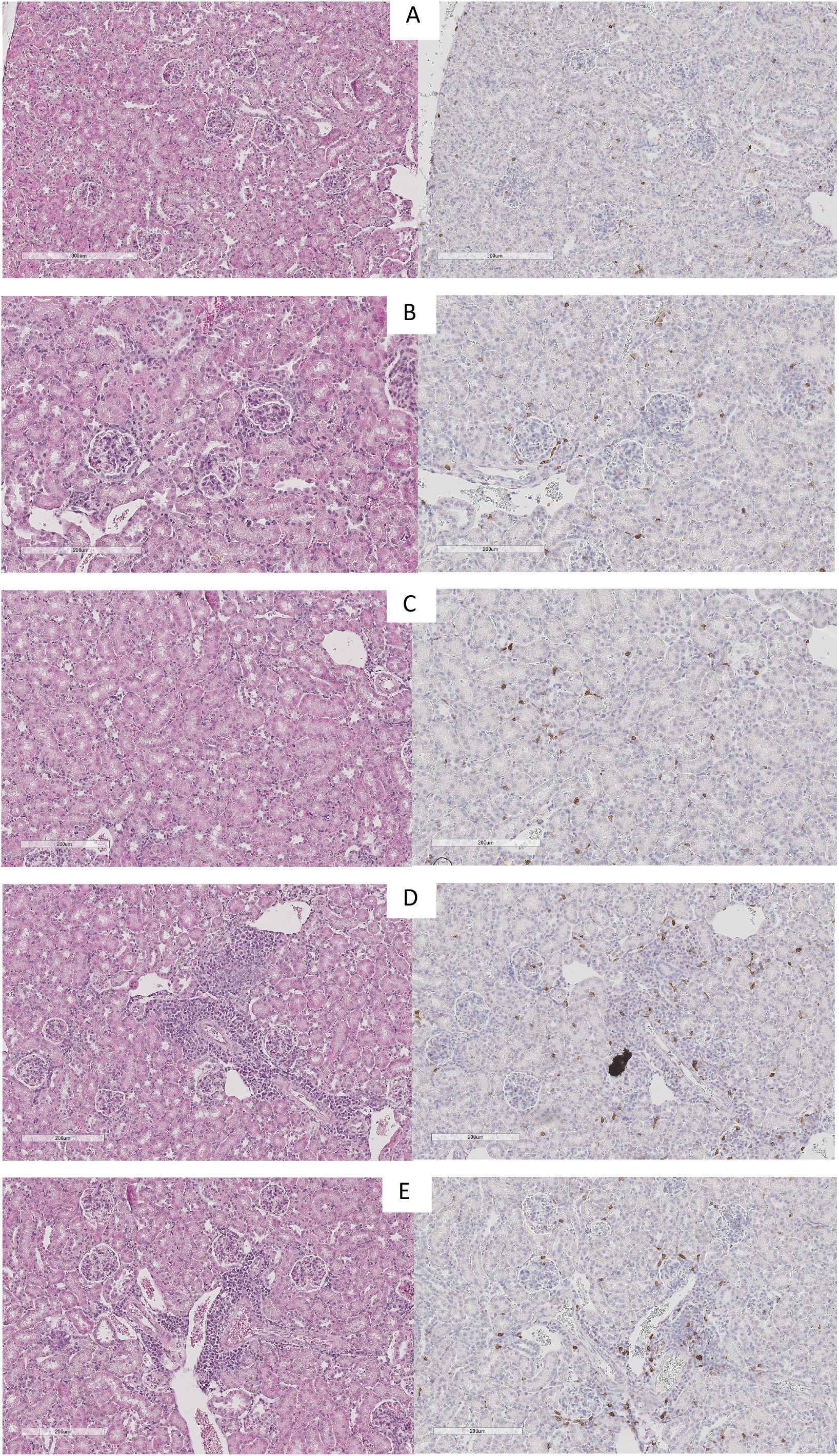
Localization of CD8^+^ T cell infiltrates in the kidney of a B6.Sle1.Sle2.Sle3 mouse. Immunohistochemical analysis of a paraffin-embedded kidney from a 10-month-old B6.Sle1.Sle2.Sle3 mouse. Adjacent sections were stained with Hematoxylin–Eosin (left panels) or anti-CD8α (right panels).

### CD4^+^ and CD8^+^ T cell infiltration assessed by flow cytometry

We next used flow cytometry on kidney cell suspensions obtained by mechanical dissociation to quantify the proportions of infiltrating CD4^+^ and CD8^+^ T cells. As shown in Fig. 4, the proportions of CD4^+^ and CD8^+^ subsets among CD3^+^ T cells remained stable across ages, maintaining a normal CD4^+^/CD8^+^ ratio of approximately 2:1. Since we had previously demonstrated that the absolute numbers of infiltrating CD8^+^ T cells increased with age in lupus mice (Fig. 2), these findings indicate a parallel age-dependent increase in CD4^+^ T cells.

**Fig. 4.**
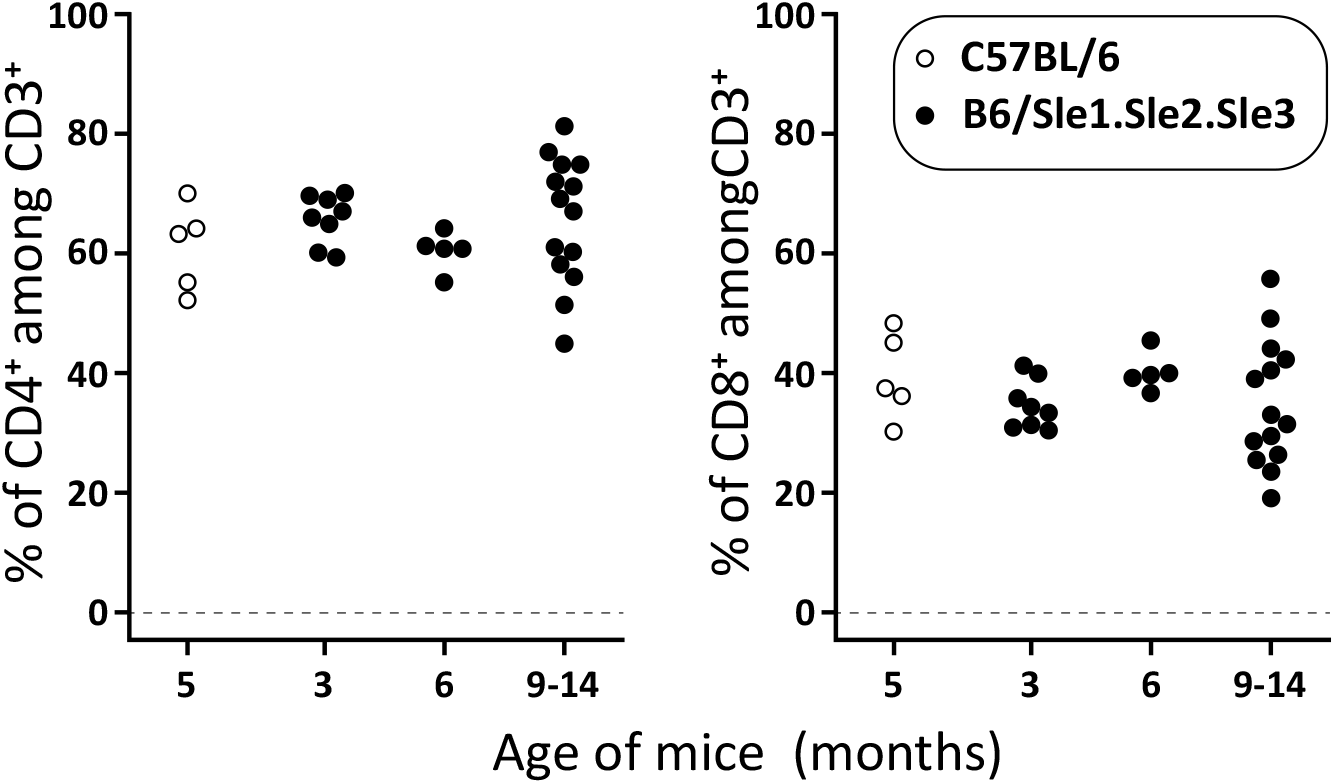
Proportions of CD4^+^ and CD8^+^ cells among T lymphocytes (CD3^+^) infiltrating the kidneys of healthy C57BL/6 and lupus B6/Sle1.Sle2.Sle3 mice. Cell suspensions prepared from kidneys (one kidney per mouse) were labeled with antibodies against CD3, CD4 and CD8, and analysed by flow cytometry.

In contrast to the study by Couzi *et al*. in human LN biopsies—where CD8^+^ T cells were reported to predominate [8]—we did not observe a predominance of CD8^+^ over CD4^+^ T cells in B6.Sle1.Sle2.Sle3 mice. Our results therefore suggest that T cell infiltration in this model is more consistent with the recruitment of circulating activated T cells (both CD4^+^ and CD8^+^) into the kidney, rather than with a local expansion or preferential activation of CD8^+^ T cells, as proposed in human LN [8-10].

Taken together, these observations suggest that the B6.Sle1.Sle2.Sle3 model may have limited relevance for dissecting the specific contribution of CD8^+^ T cells to the immunopathology of human LN.

### Gene expression profiling of lupus kidneys

To further characterize immune cell infiltration and functional pathways in the kidneys of B6.Sle1.Sle2.Sle3 lupus mice, we performed transcriptomic analyses using microarrays. Unsupervised clustering based on differentially expressed transcripts (between B6.Sle1.Sle2.Sle3 and control mice) segregated lupus mice according to age, clearly distinguishing young (3 months) from older (6–14 months) animals (Fig. 5). Heatmap analysis of 72 differentially expressed transcripts revealed several distinct expression clusters, including groups of IFN-stimulated genes that progressively increased with age. Gene Set Enrichment Analysis (GSEA) further highlighted pathways related to type 1 and type 2 IFN signaling, immune cell activation, and extracellular matrix remodeling in older lupus kidneys. Importantly, transcripts associated with B cells were strongly enriched, while T cell–associated transcripts were more heterogeneous but included cytolytic T cell genes (*Cd8a, Cd8b, Grzmb* and *Prf1*, in blue in the T cell genes panel) (Fig. 6).

**Fig. 5.**
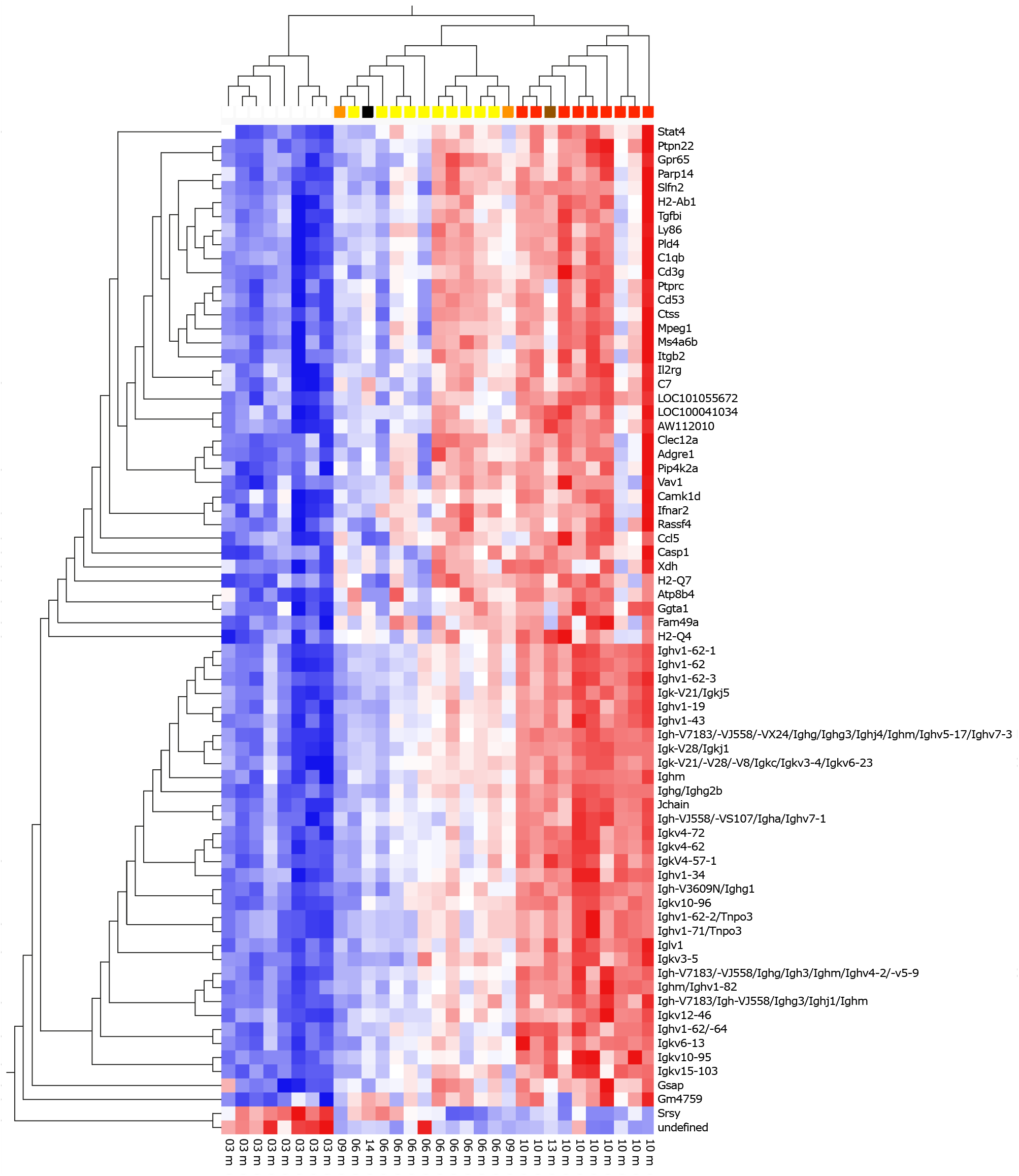
Gene expression profiling of kidneys from B6/Sle1.Sle2.Sle3 mice. Heatmap representation of supervised clustering of 72 transcripts differentially expressed in kidneys of B6/Sle1.Sle2.Sle3 lupus mice at different ages. Each column corresponds to an individual mouse, and each row to a transcript. Relative expression levels are shown in a color scale (red = upregulated, blue = downregulated). Dendrograms indicate unsupervised hierarchical clustering of samples and transcripts. Clustering segregates young (3-month-old) from older (6–14-month-old) lupus mice, consistent with age-dependent progression of renal disease. Distinct gene clusters include an IFN-inducible signature that becomes more prominent with age, as well as groups of transcripts enriched for immune cell–related pathways. These results confirm the progressive immune activation in kidneys of lupus mice and highlight similarities with human lupus nephritis.

**Fig. 6.**
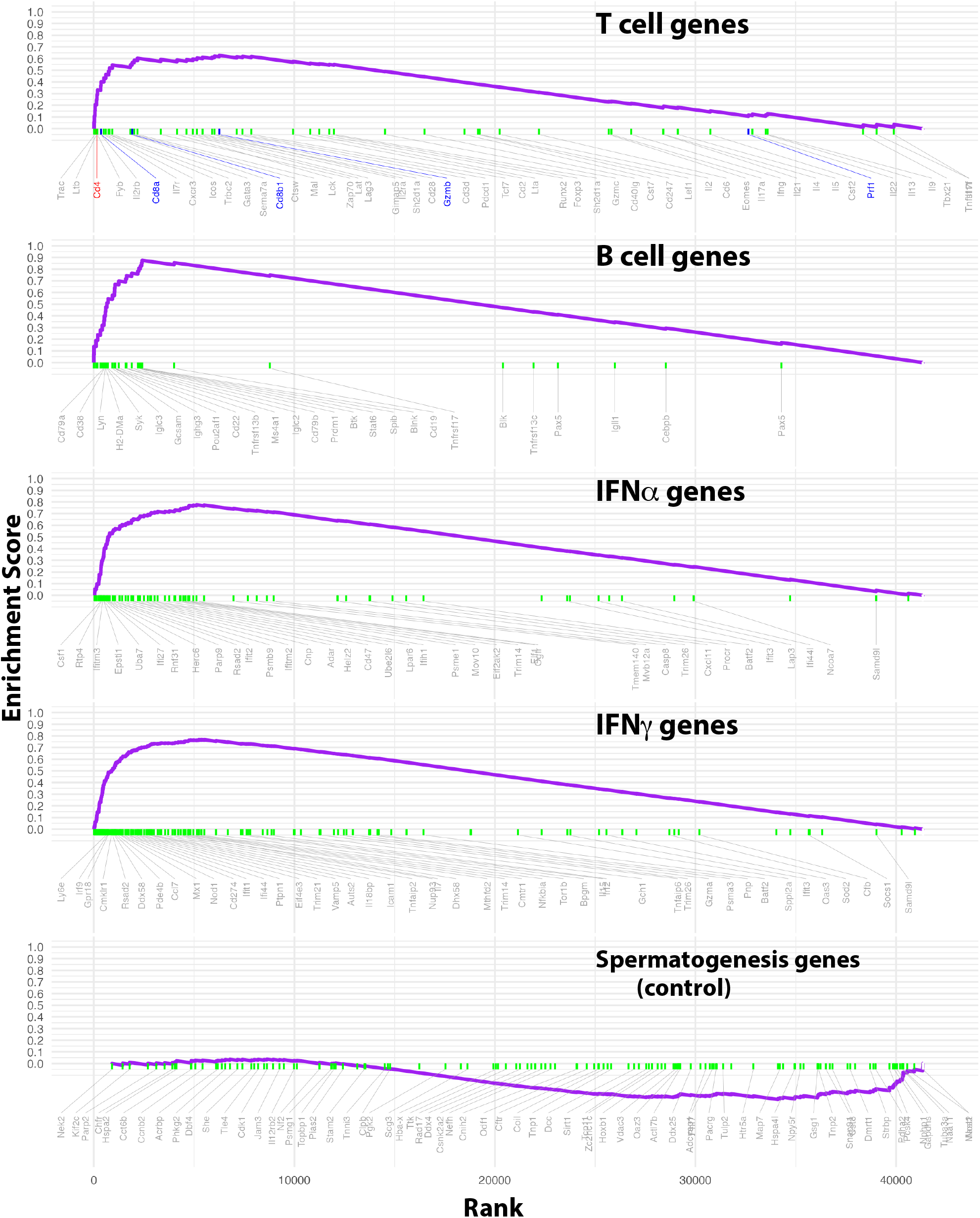
Gene Set Enrichment Analysis (GSEA) of kidney transcriptomes from B6.Sle1.Sle2.Sle3 lupus mice. Principal Component Analysis (PCA) of the microarray data showed that, as expected, age was the major source of variance in the dataset. Gene Set Enrichment Analysis (GSEA) further highlighted pathways related to type 1 and type 2 IFN signaling, immune cell activation, and extracellular matrix remodeling in older lupus kidneys. Importantly, transcripts associated with B cells were strongly enriched, while T cell– associated transcripts were more heterogeneous. The results confirmed adaptive immune infiltration and IFN-driven responses.

## Discussion

The present study demonstrates, for the first time, clear differences in CD8^+^ T cell infiltration between the kidneys of B6.Sle1.Sle2.Sle3 lupus-prone mice and those of healthy C57BL/6 controls. First, higher numbers of CD8^+^ T cells were already detectable in lupus mice at 3 months of age compared with healthy animals at any age. Second, the proportions of CD8^+^ cells progressively increased with age in lupus mice, whereas they remained consistently below 0.3% of nucleated cells in controls. Finally, CD8^+^ T cell infiltration positively correlated with NIH activity indices. Together, these findings suggest—but do not prove—a potential contribution of CD8^+^ T lymphocytes to the pathogenesis of LN in this model.

Our observations in mice are consistent with reports of CD8^+^ T cell infiltration in the kidneys of patients with LN [4–10], including the correlation between CD8^+^ cell density and histological activity [7]. However, unlike in several human studies where CD8^+^ T cells predominated among infiltrating lymphocytes [4,7], B6.Sle1.Sle2.Sle3 mice did not exhibit such predominance. Moreover, we did not identify a consistent anatomical localization of CD8^+^ T cells, in contrast to the well-defined periglomerular CD8^+^ “caps” containing oligoclonal TCR repertoires described by Winchester et al. [8].

In our view, the strongest evidence supporting a pathogenic role for CD8^+^ T cells in human LN is their frequent predominance over CD4^+^ T cells within kidney infiltrates. Whereas the physiological CD4/CD8 ratio in blood is approximately 2:1, Couzi et al. reported kidney CD4/CD8 ratios below 0.5 [7]. Such predominance may reflect either selective recruitment of CD8^+^ T cells into the kidney or local proliferation following antigen recognition. Because general mechanisms of T cell recruitment (e.g., for activated or memory subsets) do not discriminate between CD4^+^ and CD8^+^ cells, selective recruitment alone seems unlikely. A more plausible explanation is sustained, local activation of CD8^+^ T cells by renal antigens presented on HLA class I molecules. This scenario would imply antigen-driven expansion of CD8^+^ clones specifically recognizing kidney-restricted targets. Demonstrating such a role would require the identification of cognate antigens, a technically challenging task but one that could build on strategies developed for tumor antigen discovery, such as cDNA library screening. This approach would be more feasible in murine models than with the limited tissue available from patient biopsies. However, the present data indicate that the B6.Sle1.Sle2.Sle3 model does not fully recapitulate the CD8^+^ T cell biology observed in human LN. A plausible explanation would be that in SLE patients a mild reactivation of the ubiquitous BK polyomavirus (BKPyV), favoured by immunosuppressive therapies, stimulates a kidney-restriced anti-BK virus cytolytic T cell response. BKPyV does not infect mice and mouse viruses are less likely to be reactivated in the B6.Sle1.Sle2.Sle3 mice in the absence of immunosuppression.

Nevertheless, results from our group support the overall relevance of the B6.Sle1.Sle2.Sle3 model to lupus pathophysiology. In patients with LN, p16^INK4a^ senescent renal cells have been associated with both fibrosis and CD8^+^ T cell infiltration, and glomeruli with high p16 ^INK4a^ cell burden showed increased periglomerular CD8^+^ cells [10]. We observed similar features in B6.Sle1.Sle2.Sle3 mice, including renal p16 ^INK4a^ cells, their association with disease severity, and correlation with CD8^+^ infiltration [13]. These findings suggest a potential link between senescence, fibrosis, and CD8^+^ T cell–mediated pathology. The next critical step will be to identify the antigens recognized by renal CD8^+^ T cells.

Finally, recent therapeutic advances highlight the need to reassess the role of CD8^+^ T cells in SLE. Striking clinical efficacy has been reported in refractory SLE following anti-CD19 CAR T cell therapy [20]. If long-lasting remission can be achieved by depleting B cells alone, without targeting CD8^+^ T cells, the contribution of the latter to SLE pathogenesis may be less important than previously assumed.

